# COUP-TFI regulates diameter of the axon initial segment in the mammalian neocortex

**DOI:** 10.1101/2020.10.07.329284

**Authors:** Xuanyuan Wu, Haixiang Li, Jiechang Huang, Cheng Xiao, Shuijin He

**Affiliations:** School of Life Science and Technology, ShanghaiTech University, 393 Middle Huaxia Road, Pudong New District, Shanghai, 201210, CN; Institute of Neuroscience, Shanghai Institutes for Biological Sciences, Chinese Academy of Sciences, Shanghai, China; University of Chinese Academy of Sciences, Beijing, China; School of anesthesiology, Xuzhou Medical University, 209 Tongshan Road, KJL-D423, Xuzhou, Jiangsu Province, 221004, CN

## Abstract

The axon initial segment is a specialized structure that controls neuronal excitability by generating action potentials. Currently, AIS plasticity with regard to changes in length and location in response to neural activity has been extensively investigated, but how AIS diameter is regulated remains elusive. Here we report that COUP-TFI is an essential regulator of AIS diameter in both developing and adult mouse neocortex. Embryonic ablation of COUP-TFI prevented expansion of AIS diameter that occurs during postnatal development in layer II/III pyramidal cells of the mouse motor cortex, thereby leading to an impairment of action potential generation. Inactivation of COUP-TFI in adult neurons also led to reduced AIS diameter and impaired action potential generation. In contrast to different developmental stages, single-cell ablation and global ablation produced opposite effects on spontaneous network in COUP-TFI-deficient neurons. Further, mice exhibited less anxiety-like behaviors after postnatal inactivation of COUP-TFI induced by tamoxifen. Our results demonstrate that COUP-TFI is indispensable for both expansion and maintenance of AIS diameter and that a change in AIS diameter fine-tunes synaptic inputs through a metaplasticity mechanism in the adult neocortex.

## Introduction

The axon initial segment is a small specialized region that contains a high density of voltage-gated sodium and potassium channels, promoting action potential (AP) generation in neurons (Hu et al., 2009). These ion channels are clustered by a master organizing protein AnkG, and then βIV-spectrin anchors AnkG to the actin cytoskeleton (Huang and Rasband, 2018). Superresolution imaging reveals a periodic structure of the AIS in which the cytoskeletal proteins form rings spaced ∼190 nm apart (Leterrier et al., 2015; Xu et al., 2013). The AIS begins to assemble shortly after a neuron is born at the embryonic stage (Le Bras et al., 2014). AIS assembly begins with restriction of AnkG to the proximal part of the axon and thereafter AnkG recruits other AIS proteins to this region (Galiano et al., 2012; Jenkins and Bennett, 2001; Zhou et al., 1998). Several lines of evidence showed that in the absence of AnkG the AIS fails to recruit ion channels and the major cytoskeletal proteins to the proximal axon (Hedstrom et al., 2007; Sobotzik et al., 2009; Zhou et al., 1998).

The AIS varies in location and length along the axon across different subtypes of neurons, both of which are crucial for modulating neural excitability and controlling the temporal precision of spike coding (Grubb and Burrone, 2010; Kuba et al., 2006; Kuba et al., 2010; Lazarov et al., 2018). Recently, AIS plasticity has been well documented as changes in the length and location in response to neural activity *in vitro* and *in vivo* (Evans et al., 2013; Grubb and Burrone, 2010; Kuba et al., 2010). For instance, AIS shortening during development is prevented by deprivation of sensory inputs (Gutzmann et al., 2014; Kuba et al., 2010). A further mechanistic study demonstrates that the Ca^2+^-calmodulin-dependent calcineurin signaling is involved in AIS plasticity (Evans et al., 2013), which is initiated by calcium influx through L-type calcium channels. Over the past decades, numerous studies focus on how a neuron controls its AIS length and location; however, AIS diameter has received little attentions from scientists. A previous study showed that AIS diameter is enlarged in the spinal cord of human patients with amyotrophic lateral sclerosis (ALS) (Sasaki and Maruyama, 1992). AIS diameter varies across different neurons and shows a positive relationship with the size of soma (Sasaki and Maruyama, 1992), but little is known about the factors that control AIS diameter in mature neurons.

In this study, we firstly found that Emx1-Cre-dependent conditional ablation of COUP-TFI (Chicken Ovalbumin Upstream Promoter-Transcription Promoter 1, also known as Nr2f1) resulted in reduced AIS diameter, a more positive AP threshold and reduced frequency of AP-dependent spontaneous postsynaptic currents (sPSCs) in cortical pyramidal cells. We then deleted COUP-TFI in single neurons by injection of viruses carrying iCre into the lateral ventricle of embryos (retrovirus) or into the adult neocortex (adeno-associated virus). Compared with the global knockout, loss of COUP-TFI in single neurons produced similar effects on AIS diameter and AP properties, but increased the frequency of sPSCs and miniature excitatory postsynaptic currents (mEPSCs) as well as the density of spines. In contrast, overexpression of COUP-TFI in single neurons led to increased AIS diameter, a more negative AP threshold and reduced AP-synaptic transmission. Furthermore, postnatal ablation of COUP-TFI by tamoxifen (TMX) reduced anxiety-like behaviors in mice, indicating a correlation between AIS diameter and cognitive function. Taken together, these data suggest that AIS diameter is capable of being regulated by COUP-TFI and tightly linked with metaplasticity in the adult mammalian neocortex.

## Results

### COUP-TFI regulates axon initial segment diameter during postnatal development

To investigate development of AIS diameter, we immunostained AnkG, a general marker for the AIS, in the mouse neocortex. We analyzed diameter of the AIS at the start (closest to soma) and middle points in pyramidal neurons within the lay II/III of the motor cortex, and found that AIS diameter was progressively increasing during normal postnatal development. Accidently, we found that AIS diameter significantly reduced in cortical pyramidal cells from Emx1-Cre; COUP-TFI flox/flox (COUP-TFI fl/fl) mice (referred to COUP-TFI cKO), compared with COUP-TFI fl/fl mice (referred to controls) (Suppl. Fig. 1). This decrease occurred after P7 (Suppl. Fig. 1A-C), when the AIS completes its assembly but yet undergoes diameter expansion (Suppl. Fig. 1B-C), suggesting that COUP-TFI is crucial for AIS diameter expansion instead of assembly (Hedstrom et al., 2007). In contrast, ablation of COUP-TFI in the hippocampus caused the bending and misdirection of the AIS rather than affecting the diameter in CA1 pyramidal cells (Suppl. Fig. 2), reminiscent of the bending axon (Armentano et al., 2006). We then restricted our analyses to the layer II/III of the motor cortex for this study because this region preserves in COUP-TFI cKO mice. Immunostaining of another AIS marker βIV-spectrin confirmed this finding from AnkG staining in the neocortex (Suppl. Fig. 1H-J). Immunoblot revealed no change in expression of the AIS proteins, AnkG, βIV-spectrin and Nav1.6, between cKO mice and controls (Suppl. Fig. 1K, L), suggesting that reduced AIS diameter is not due to a decrease in protein expression.

Numerous studies previously reported that COUP-TFI plays important roles in cortical arealization/lamination, neuronal migration, and neural circuit formation as well as specification of neuronal identity (Bertacchi et al., 2019). These raise the possibility that neurons in the same cortical region of controls and cKO mice might belong to different subtypes of pyramidal cells, which could confound interpretation of these data related to alterations in AIS diameter. To circumvent this problem, we ablated COUP-TFI in single neurons by delivery of low-titer retroviruses expressing iCre fused with enhanced GFP (EGFP) into the lateral ventricle of embryos from pregnant COUP-TF fl/fl mice at embryonic day 14.5 (Fig. 1A), when the majority of new-born neurons migrate into the superficial layer (Layer II-IV). Retroviruses expressing EGFP only serves as a control. Notably, COUP-TFI was efficiently deleted in iCre-expressing neurons (93%; 27 of 29, Fig. 1B, C). Consistent with results from cKO mice, we observed a significant reduction of AIS diameter following loss of COUP-TFI in single cells (Fig. 1D, E). In contrast, we did not see a change in AIS diameter at the distal end point (Fig. 1E), indicating that axonal diameter is not affected by deletion of COUP-TFI. The length of AIS was not changed after loss of COUP-TFI (Fig. 1F). Since cell size and AIS location appear to have impacts on AIS diameter (Sasaki and Maruyama, 1992), we performed analyses of maximal cell area and the distance between the AIS and soma. We found that neither of them was changed between EGFP only and iCre-expressing neurons (Fig. 1G, H), ruling out these possibilities. Collectively, these results demonstrate that regulation of the AIS by COUP-TFI is specific to the diameter.

**Figure 1:**
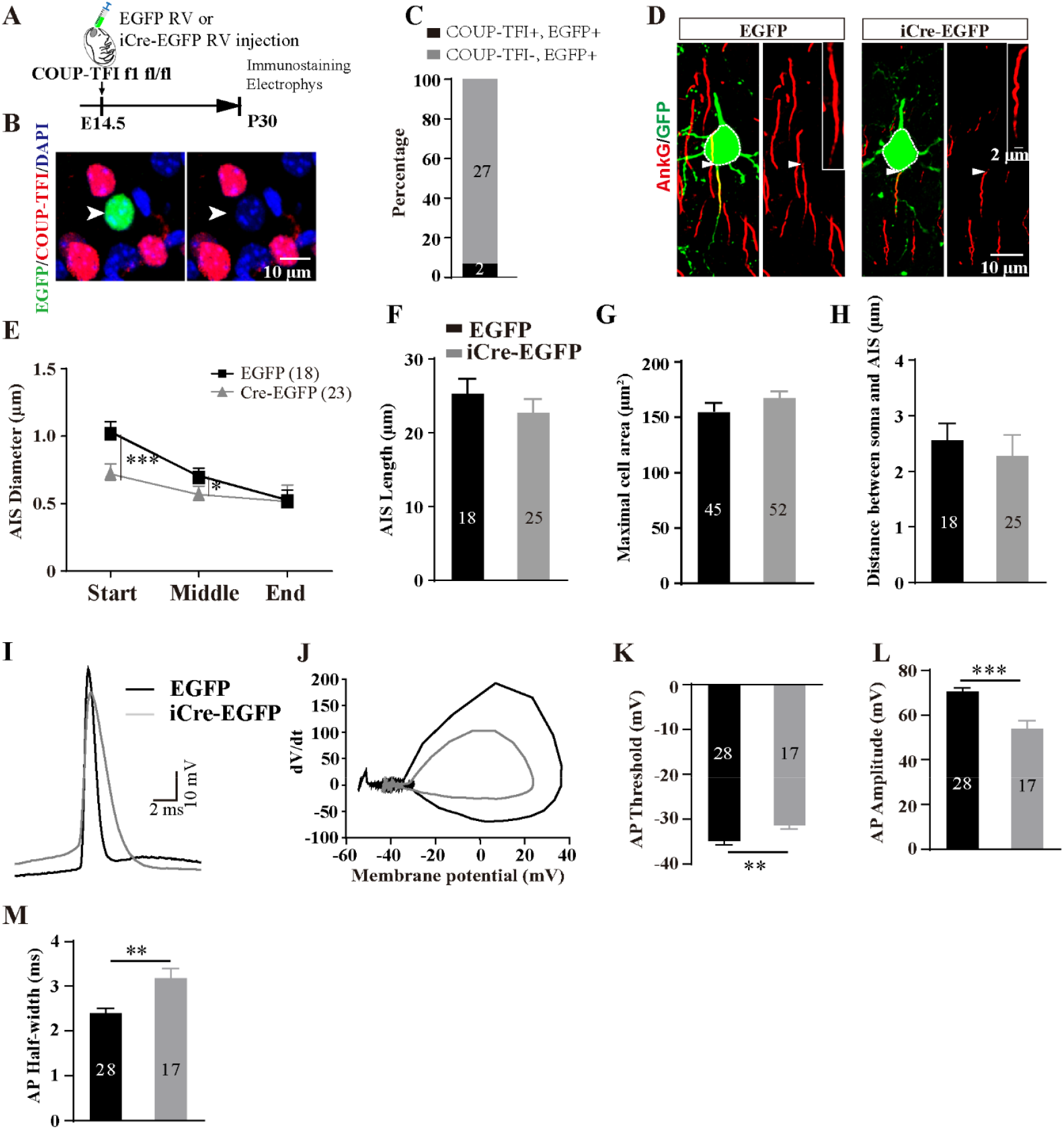
Loss of COUP-TFI in single neurons reduces AIS diameter and impairs AIS function. (A) Schematic of the experimental paradigm. Mice that received retroviruses (RV) expressing EGFP or iCre fused EGFP at embryonic day 14.5 were sacrificed at P30 for immunostaining and electrophysiological recordings. (B) Representative confocal images of co-staining of EGFP (green) and COUP-TFI (red). Arrow heads indicate an EGFP+, COUP-TFI-cell. (C) Quantification for assessing the knockout efficiency by iCre (n= 3 mice). (D) Representative images of AnkG immunostaining for EGFP-or iCre-EGFP-infected neurons in the layer II/III of the mouse neocortex. Boxed regions are the expanded AISs. Arrow heads indicate the start point of the AIS. Dash lines circle the maximal area of cells. (E) Quantification of AIS diameter at the start, middle and end points in EGFP-or iCre-EGFP-expressing neurons. (F-H) Quantification of the length of the AIS (*F*), the maximal cell area (*G*), and the distance between the AIS and the soma (*H*). (E-H: EGFP, n represents neuron number from 4 mice; Cre-EGFP, n represents neuron number from 4 mice). (I) Sample traces of the first APs were overlapped. (J) Phase-plane plots of membrane voltage and its change rate in response to current injections. (K-M) Quantification of action potential (AP) threshold (*K*), peak amplitude (*L*), and half-width (*M*). (K-M: EGFP, n represents neurons from 6 mice; Cre-EGFP, n represents neurons from 4 mice). Data are presented as mean ± SEM. *, p < 0.05; **, p < 0.01; ***, p < 0.001, student’s t-test.

To examine functional consequence of reduced AIS diameter, we performed whole-cell electrophysiological recordings of action potentials in cortical neurons. We firstly determined the relationship between AIS diameter and function. Apparently, the larger AIS diameter, the more negative AP threshold, the larger AP amplitude, and the narrower half-width (Suppl. Fig. 3). Consistent with this idea, neurons expressing iCre-EGFP displayed more positive AP threshold, smaller AP amplitude and broadened half-width, compared with controls (Fig. 1 I-M).

We next asked whether overexpression (OE) of COUP-TFI could enlarge AIS diameter. Similarly, retroviruses for expression of EGFP only or COUP-TFI-IRES-EGFP were delivered into the lateral ventricle of the embryos at E14.5. In contrast to EGFP only, AIS diameter significantly increased in COUP-TFI OE neurons (Fig. 2A-C), resulting in elevated AIS functions, including a more negative AP threshold and increased AP amplitude (Fig. 2D-H). Taken together, these results suggest that COUP-TFI regulates expansion of AIS diameter during postnatal development.

**Figure 2:**
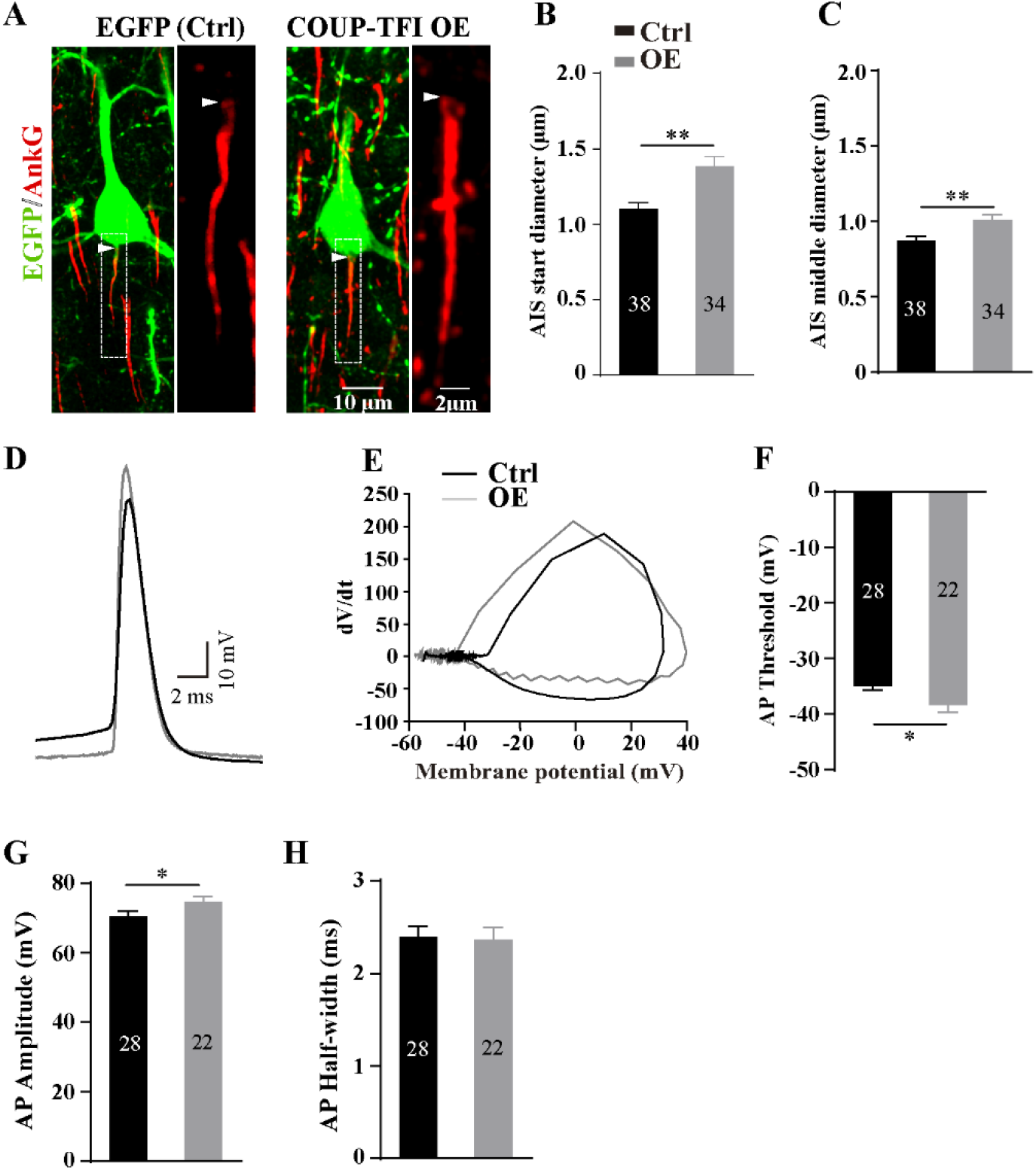
Overexpression of COUP-TFI in single neurons enlarges AIS diameter and enhances AIS function. (A) Representative confocal images of co-staining of EGFP (green) and AnkG (red). Boxes are expanded to the right panels. Arrow heads indicate the start point of the AIS. (B, C) Quantification of AIS diameter at the start (*B*) and middle (*C*) points in neurons expressing EGFP- or COUP-TFI-IRES-EGFP. (B-C: EGFP, n represents neurons from 4 mice; COUP-TFI OE, n represents neurons from 4 mice). (D) Sample traces of the first APs were overlapped. (E) Phase-plane plots of membrane voltage and its change rate in response to current injection. (F-H) Quantification of AP threshold (*F*), peak amplitude (*G*), and half-width (*H*) for the first AP evoked. (F-H: EGFP, n represents neurons from 6 mice; COUP-TFI OE, n represents neurons from 8 mice). Data are presented as mean ± SEM. *, p < 0.05; **, p < 0.01, student’s t-test.

### COUP-TFI is required for maintenance of AIS diameter in the adult neocortex

Could AIS diameter be regulated in mature cortical neurons? To study this question, we performed injection of adeno-associated viruses carrying EGFP only or iCre-P2A-EGFP into the layer II/III of the motor cortex of COUP-TFI fl/fl mice at P30 (Fig. 3A). We found that AIS diameter significantly reduced (Fig. 3B-D), accompanying by a more positive AP threshold and a decrease in AP amplitude (Fig. 3E-I), in iCre-P2A-EGFP-expressing neurons, compared with EGFP only, indicating a tight link between AIS diameter and neuronal excitability. These results suggest that COUP-TFI is required for maintenance of AIS diameter in adult cortical neurons.

**Figure 3:**
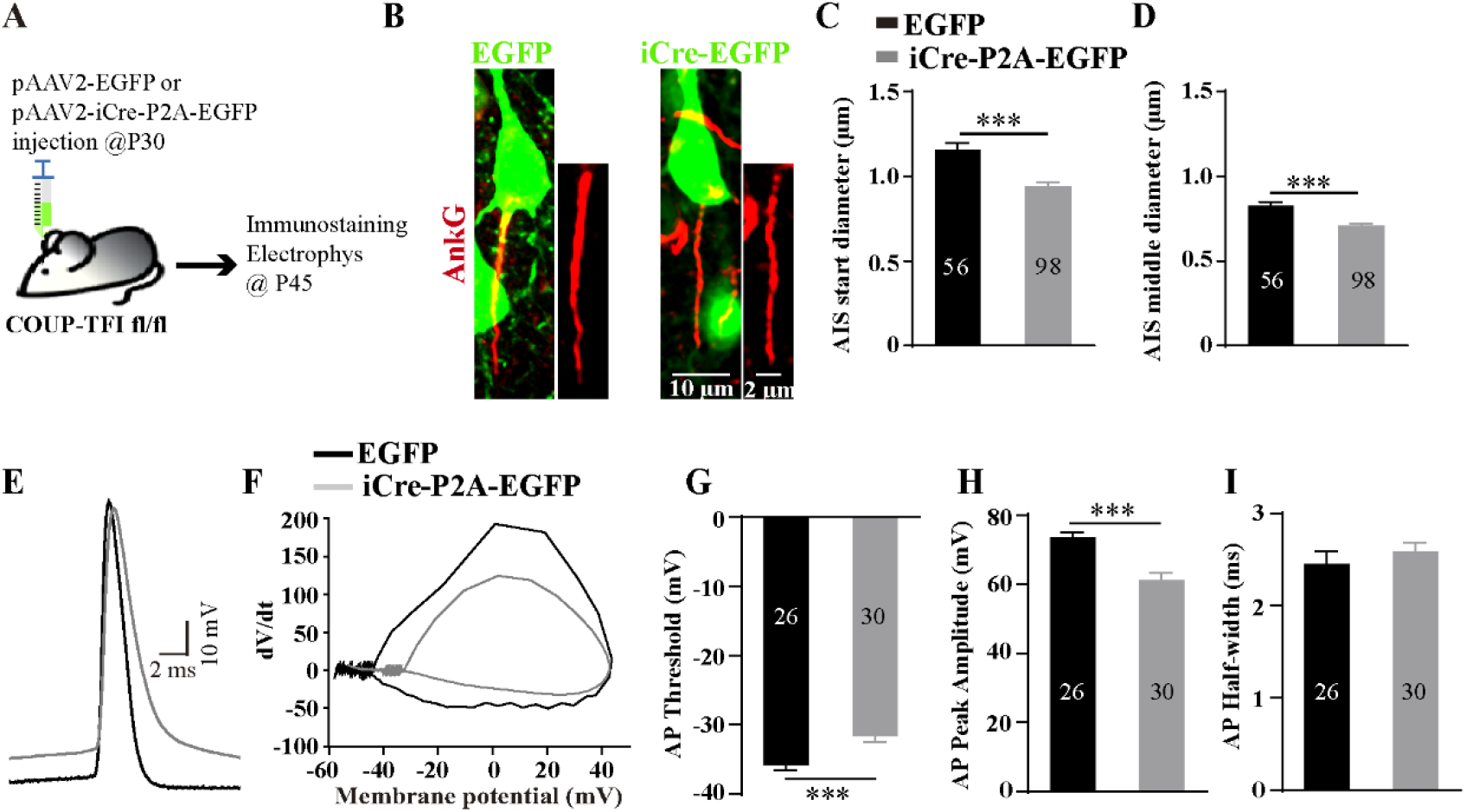
COUP-TFI is required for maintenance of AIS diameter in adult pyramidal cells. (A) Schematic of the experimental paradigm. Adeno-associated viruses that express EGFP or iCre-P2A-EGFP were injected into the lay II/III of the mouse neocortex at P30 and then mice were sacrificed ∼15 days later for further experiments. (B) Representative confocal images of AnkG for these groups. AISs are expanded to the right panels. (C, D) Quantification of AIS diameter at the start (*C*) and middle (*D*) points in neurons expressing EGFP only or iCre-P2A-EGFP. (C-D: EGFP, n represents neurons from 4 mice; iCre-P2A-EGFP, n represents neurons from 6 mice). (E) Sample traces of the first APs were overlapped. (F) Phase-plane plots of membrane voltage and its change rate in response to current injections. (G-I) Quantification of AP threshold (*G*), peak amplitude (*H*), and half-width (*I*) for the first AP initiated. (G-I: EGFP, n represents neurons from 4 mice; iCre-P2A-EGFP, n represents neurons from 5 mice) Data are presented as mean ± SEM. ***, p < 0.001, student’s t-test.

### Tuning of synaptic plasticity by AIS diameter

Neuronal excitability is fine-tuned through altering AIS location and length in response to synaptic inputs or ongoing neuronal activities (Grubb and Burrone, 2010; Kuba et al., 2010). Conversely, synaptic inputs can be re-scaled by chronically altered neuronal excitability for compensating neuronal outputs (Burrone and Murthy, 2003; Turrigiano and Nelson, 2000). Based on our results showing an association between AIS diameter and neuronal excitability, we reasoned that synaptic inputs could be fine-tuned by altering AIS diameter. We examined the spontaneous network on COUP-TFI-deficient neurons from cKO mice and from mice that received retrovirus injections at the embryonic stage. We found that the frequency of AP-dependent sPSCs evidently reduced in COUP-TFI cKO mice (Fig. 4A-D), but no changes in AP-independent miniature PSCs (mPSCs) (Fig. 4E-G), compared with controls. Conversely, sPSC frequency significantly increased in single iCre-EGFP-expression neurons in contrast to EGFP only (Fig. 4J-L). Interestingly, overexpression of COUP-TFI in single neurons produced effects on sPSCs similar to those of COUP-TFI cKO mice, but contrary to those of single COUP-TFI-deficient neurons (Fig. 4J-L). Thus, these results suggest that these effects on spontaneous network in single neurons likely results from homeostatic plasticity rather than direct genetic regulation of synaptic inputs by COUP-TFI; that is to say, the gain of synaptic inputs is scaled to compensate a reduction in neuronal excitability that results from decreased AIS diameter.

**Figure 4:**
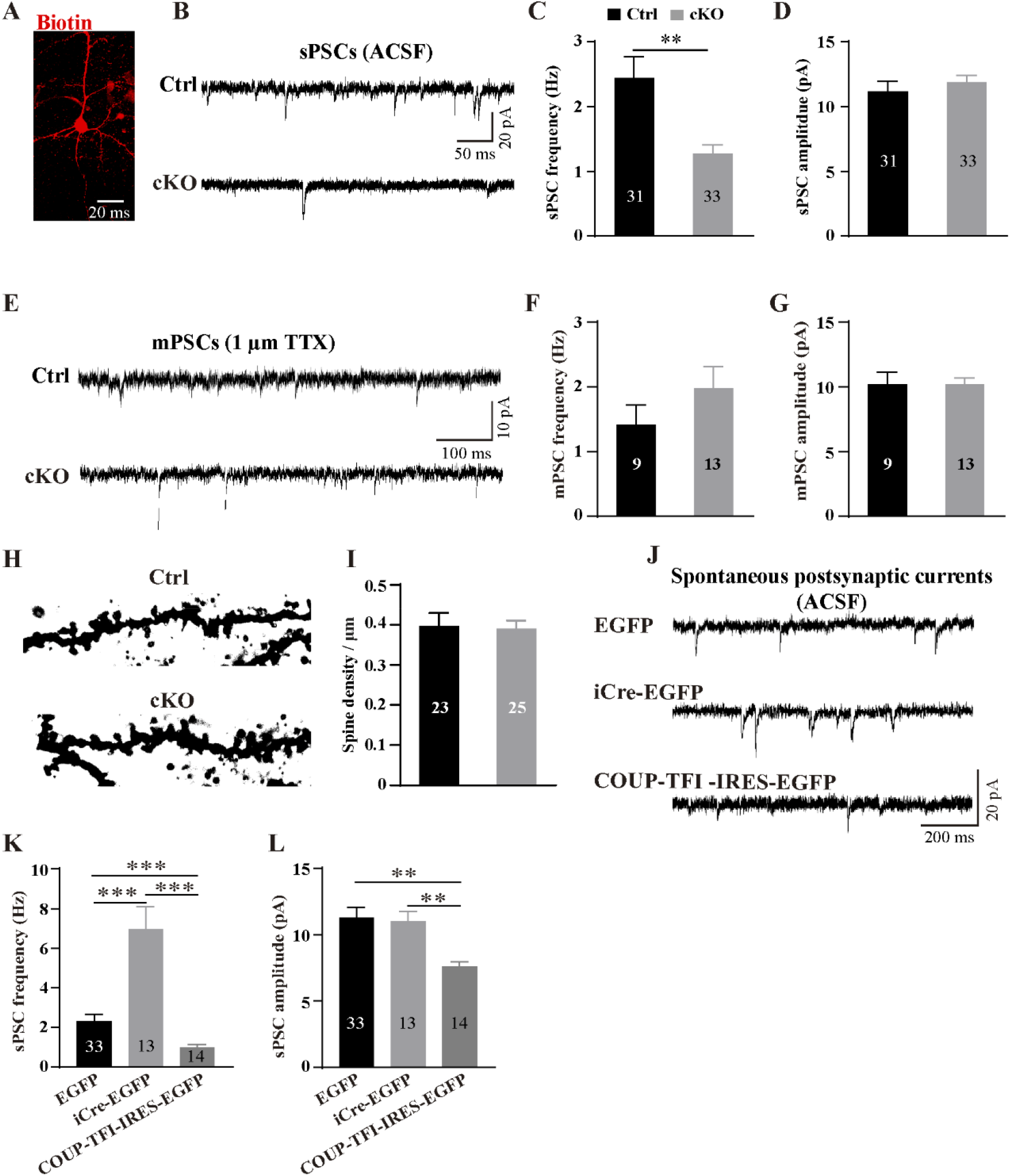
Loss of COUP-TFI at the single-cell level and the global level produces opposite effects on the receiving spontaneous network. (A) Representative confocal images of a pyramidal neuron filled with neurobiotin from a control. (B) Sample traces of spontaneous postsynaptic currents (sPSCs) from P30 mice. (C, D) Quantification of the frequency and amplitude of sPSCs in cortical pyramidal cells. (C-D: Ctrl, n represents neurons from 5 mice; cKO, n represents neurons from 7 mice). (E-G) Sample traces (*E*) and quantification of the frequency (*F*) and amplitude (*G*) of miniature postsynaptic currents (mPSCs) recorded in the ACSF containing 1 µm TTX. (F-G: Ctrl, n represents neurons from 3 mice; cKO, n represents neurons from 3 mice). (H, I) Representative images (*H*) and quantification (*I*) of spines in the pyramidal cells in the neocortices from control and cKO mice. (I: Ctrl, n represents neurons from 3 mice; cKO, n represents neurons from 3 mice). (J-L) Sample traces (*J*) and quantification of the frequency (*K*) and amplitude (*L*) of sPSCs recorded in the single EGFP-positive cortical pyramidal neurons from mice that received retroviruses expressing EGFP only, iCre-EGFP or COUP-TFI-IRES-EGFP at the embryonic stage. (K and L: EGFP, n represents neurons from 6 mice; iCre-EGFP, n represents neurons from 4 mice; COUP-TFI-IRES-EGFP, n represents neurons from 8 mice). Data are presented as mean ± SEM. **, p < 0.01; ***, p < 0.001. P value in (C) is determined by student’s t-test. P values in (K) and (L) are calculated using One-way ANOVA with *post ho*c Bonferroni’s tests.

We then examined synaptic plasticity in the adult mouse brain. Similarly, Spsc frequency significantly increased in single COUP-TFI-deficient neurons from mice that received AAV injection at P30, compared with controls (Fig. 5A-C). This increase is owing to upregulated excitatory synaptic transmission because we found increases in the frequency of mEPSCs (Fig. 5D-F) and the density of spines (Fig. 5J-K), but no significant changes in miniature inhibitory postsynaptic currents (mIPSCs) (Fig. 5G-I). Notably, the spine density was not altered in COUP-TFI cKO mice (Fig. 4H, I). Taken together, these data suggest that AIS diameter fine-tunes synaptic inputs on neurons in the adult neocortex, crucial for maintenance of homeostasis.

**Figure 5:**
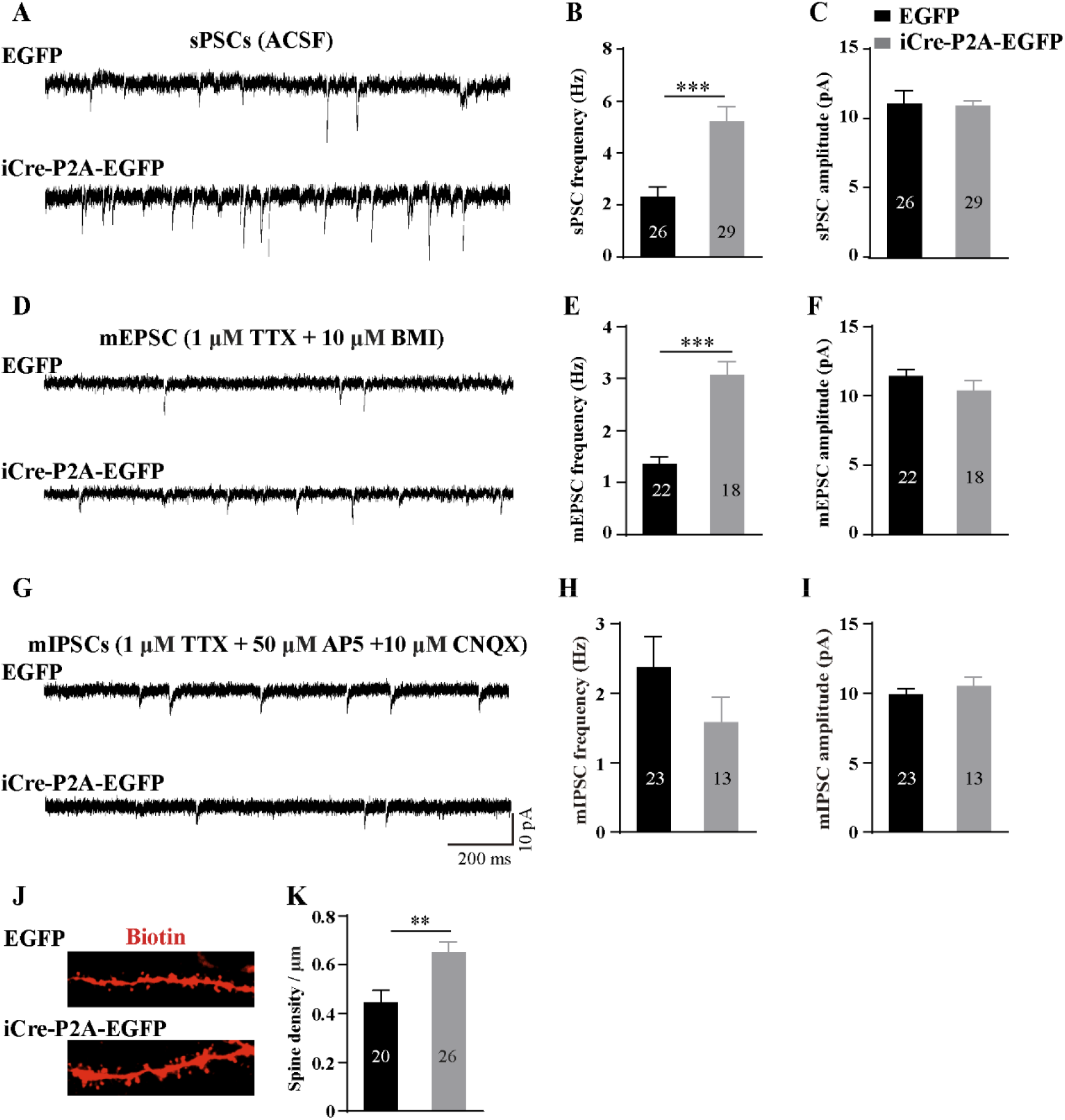
Loss of COUP-TFI in single adult neurons impairs excitatory synaptic transmission. (A-I) Sample traces and quantification of sPSCs (*A-C*), miniature excitatory postsynaptic currents (mEPSCs) (*D-F*), and miniature inhibitory postsynaptic currents (mIPSCs) (*G-I*) from mice that received injection of AAVs at P30. (J, K) Representative images (*J*) and quantification (*K*) of spines in the pyramidal cells recorded with the pipette solution containing biotin. (B-K: EGFP, n represents neurons from 4 mice; iCre-P2A-EGFP, n represents neurons from 5 mice). Data are presented as mean ± SEM. **, p < 0.01; ***, p < 0.001, student’s t-test.

### Postnatal deletion of COUP-TFI alleviates anxiety-like behaviors in mice

To further investigate possible roles of AIS diameter in brain functions, we performed intraperitoneal injection of TMX into COUP-TF fl/fl^Emx1-CreER^ mice at P15 (Fig. 6A), when neuronal migration, axonal projection and cortical patterning are completed. This TMX injection time would minimize effects of alterations in these developmental processes on brain functions. Immunostaining confirmed ablation of COUP-TFI in COUP-TF fl/fl^Emx1-CreER^ mice (Suppl. Fig. 4B). In agreement with the results from AAV experiments, AIS diameter evidently decreased in cortical neurons from COUP-TF fl/fl^Emx1-CreER^ mice, compared with COUP-TF1fl/fl (Suppl. Fig. 4C-E). We then performed the open field and elevated plus maze tests to assess mouse anxiety approximately 10 weeks after TMX injection (∼12-13 weeks old), respectively. Importantly, we found that in contrast to COUP-TF1 fl/fl mice, COUP-TF fl/fl^Emx1-CreER^ mice irrespective of gender spent more time in the center and travelled more distance in the open field task (Fig. 6B-D). Likewise, these mice spent more time in the open arm of the elevated open maze, indicating reduced anxiety-like behaviors (Fig. 6E-G). Taken together, these data suggest that postnatal deletion of COUP-TFI alleviates mouse anxiety, which correlates with reduced AIS diameter.

**Figure 6:**
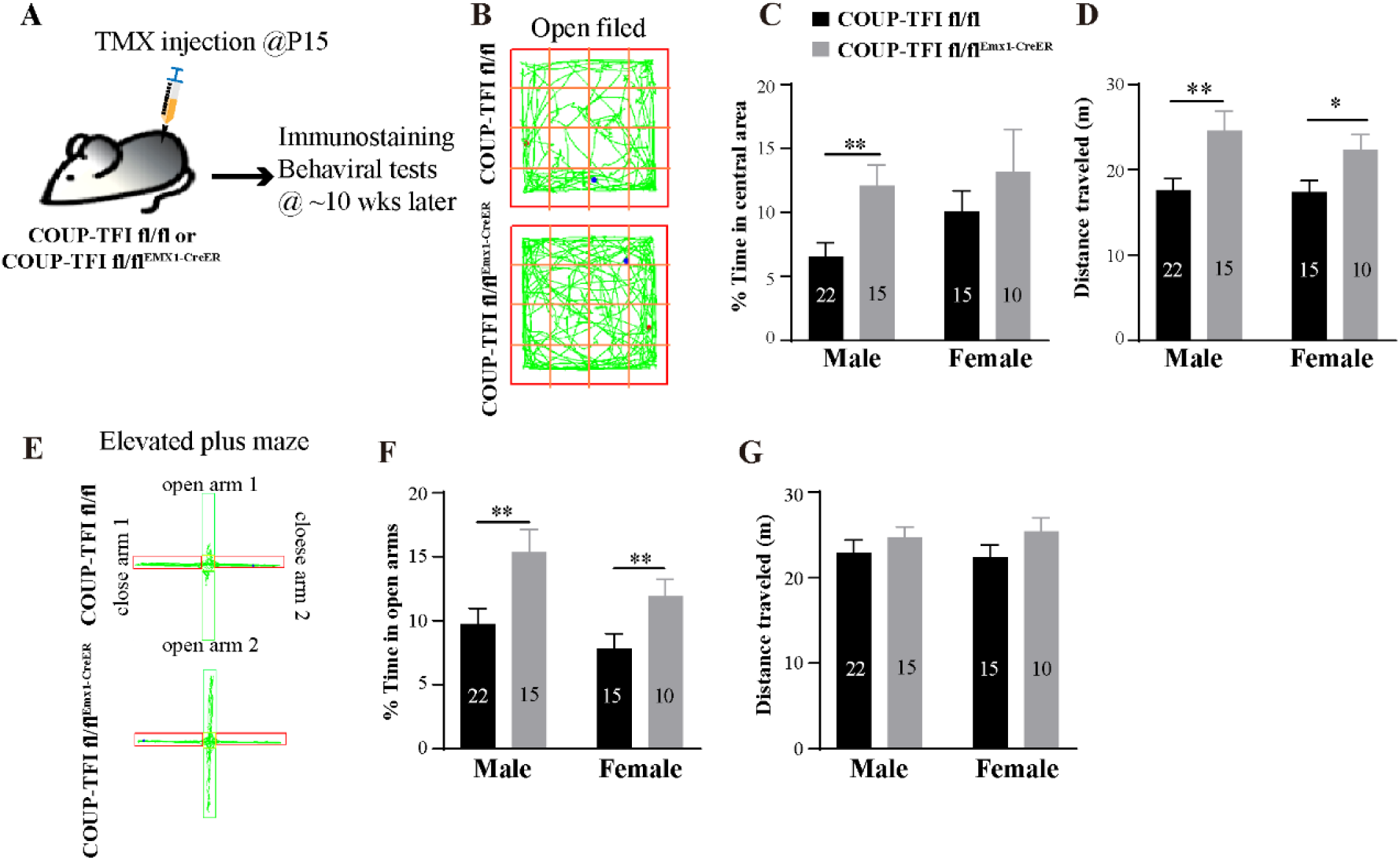
Postnatal ablation of COUP-TFI reduces anxiety-like behaviors in mice. (A) Schematic of the experimental paradigm for behavioral tests. Mice received intraperitoneal injection of tamoxifen at P15 and behavioral tests were performed ∼10 weeks later. (B) Sample trajectory of mouse activity in the open field. (C, D) Quantification of time spent in the center of the open field (C) and travel distance (D) for the male and female mice, respectively. (E-G) Sample trajectory (E) of mouse activity and quantification of time spent in the open arm (C) and travel distance (D) in the elevated plus maze for the male and female mice, respectively. N represents mouse number assessed. *, p < 0.05; **, p < 0.01, student’s t-test.

## Discussion

In this study, we found that ablation of COUP-TFI significantly led to a reduction of the diameter but no change in the length of the AIS in layer II/III pyramidal cells of the mouse motor cortex. In contrast to the cortex, we found unchanged AIS diameter but misdirection in hippocampal neurons from COUP-TFI cKO mice, compared with controls. A most recent study reported that both AIS diameter and length significantly decreased in layer V pyramidal cells of COUP-TFI cKO mouse somatosensory cortex compared with controls (Del Pino et al., 2020), suggesting that COUP-TFI might produce region- and/or layer-specific effects on the AIS. Moreover, we found that COUP-TFI is required for maintenance of AIS diameter in mature neurons and AIS diameter of a neuron is crucial for fine-tuning its synaptic inputs.

By manipulating COUP-TFI expression in single cells, we found that ablation drastically increased while overexpression significantly reduced the frequency of sPSCs in these COOUP-TFI-deficient neurons. These differential effects on synaptic inputs parallel with changes in their AIS diameters. In contrast to single-cell deletion, global ablation of COUP-TFI reduced sPSC frequency, suggesting that increased sPSC frequency in single neurons is unlikely due to genetic effects of COUP-TFI. In support of this notion, the complexity of dendritic morphology is reduced after global ablation of COUP-TFI (Alfano et al., 2011). Thus, these bidirectional effects of COUP-TFI ablation and overexpression at the single-cell level are most likely due to homeostatic plasticity; that is to say, a change in AIS diameter produced compensating effects for altered neuronal excitability by tuning-in synaptic inputs, therefore maintaining neural homeostasis.

Although COUP-TFI has multi-faceted roles in brain development (Bertacchi et al., 2019), the role of COUP-TFI in regulation of AIS diameter in mature neurons provides us an opportunity to study the role of AIS diameter in brain functions. Indeed, we found that anxiety-like behaviors significantly reduced in TMX-induced cKO mice at P15, compared with controls. Overall, these results from TMX-induced postnatal cKO mice are in agreement with previous findings from embryonic cKO mice (Contesse et al., 2019; Flore et al., 2017). These findings suggest a correlation between AIS diameter and brain functions. At present, although early developmental processes such as cell migration, cortical arealization/lamination and axon guidance are unlikely involved in altered brain functions, we cannot rule out other possibilities that COUP-TFI may have other unknown cellular functions.

In summary, results from this study provide important insights into understanding of how a neuron maintains its AIS diameter and how AIS diameter fine-tunes synaptic inputs in the adult brain. Further investigations are needed to elucidate mechanisms responsible for controlling AIS diameter in the neocortex. Human patients carrying haploinsufficient COUP-TFI exhibited intellectual disability; therefore, it will be interested to test whether AIS diameter contributes to intellectual ability.

## Materials and Methods

### Animals

COUP-TFI flox/flox (fl/fl) C57BL/6 mice (a kind gift from Dr. Ke Tang) were crossed with Emx1-Cre knock-in C57BL/6 mice (a kind gift from Dr. Ke Tang, Mouse Genome Informatics ID: 1928281) or Emx1-CreER knock-in C57BL/6 mice (a kind gift from Dr. Yun-Chun Yu, Fudan University) to generate an Emx1-Cre, COUP-TFI fl/fl mouse line (cKO) or Emx1-creER, COUP-TFI fl/fl mouse (tamoxifen-indeced cKO) line. All mice were maintained under standard conditions of 22 ± 2°C, 50±10% relative humidity and 12-hours (hrs) light/dark cycle, with food and water available. All experiments were carried out in accordance with the animal experimental protocols approved by the Animal Care and Use Committee of the ShanghaiTech University.

### Tamoxifen injection

For conditional knock out of COUP-TFI in the Emx1-creER; COUP-TFI fl/fl mouse line, 50 mg/ml Tamoxifen (TMX, Sigma Cat #: T5648) was diluted in corn oil and ethyl alcohol mixed buffer (corn oil:ethyl alcohol 19:1). TMX solution (250 mg/kg) was intraperitoneally injected to mice at P15. Immunostaining were performed ∼4 weeks after TMX injection. Behavioral tests were performed ∼10 weeks after TMX.

### Golgi-Cox staining

Adult Emx1-Cre, COUP-TFI fl/fl mice and WT mice were used to Golgi-Cox staining. Mice were anesthetized by chloral hydrate and transcardially perfused with 0.1 M phosphate buffered saline (PBS) and then quickly washed with ddH2O for removing blood from the surface. Golgi staining was performed according to the standard protocol provided with FD Rapid GolgStain™ Kit (Cat #: PK401, FD NueroTechnologies, INC). Briefly, fresh brains were treated with the mix of Solution A and B (prepare it right before mouse sacrifice) in the dark at room temperature for 3 weeks, and then transferred into Solution C to be incubated for another 5 days. Afterwards, mouse brains were sectioned into coronal 200 μm slices using vibratome. Sections were washed with ddH2O and incubated in DE mixed buffer for 10 mins followed by ddH2O wash and dehydration with 75%, 95% and 100% ethanol. Slices were treated with xylene (Cas #: 1330-20-7, Greagent) and cover slipped with the DPX Mountant (Lot #: BVBH4393V, Sigma).

### Immunostaining

For brains tissue preparations, mice were anesthetized by chloral hydrate, transcardially perfused with paraformaldehyde (PFA) in 0.1 M PBS with pH 7.4 (1% PFA for AnkG or COUP-TFI staining and 4% PFA for EGFP or βIV-spectrin). Brains were removed, post-fixed for 3 hrs at 4°C., and then transferred into 20% sucrose in 0.1 M PBS at 4°C for over 24 hrs. The dehydrated brains were sectioned to 25 µm coronal sections using cryostat microtome. Immunostaining was performed as describes previously (Zhao et al., 2020). Primary antibodies were used in this study as follows: mouse anti-COUP-TFI (1:500; PP-H8132-00, PPMX), Rabbit anti-Nav1.6 (1:500; ASC009, Alomone), Mouse anti-AnkG (1:500, Santa Cruz), Chicken anti-GFP (1:1000; GFP-1020, Aves Lab), Rabbit anti-NeuN (1:1000, ABCam), Rabbit anti-βIV-spectrin (1:1000, a gift from Matthew N. Rasband)..

### Electrophysiological recordings

Electrophysiological recordings were performed as previously described (Jiang et al., 2015; Yu et al., 2012; Zhao et al., 2020). Adult mice were deeply anesthetized by isoflurane and transcardially perfused with ice-cold ACSF solution consisting of: 126 mM NaCl, 4.9 mM KCl, 1.2 mM KH2PO4, 2.4 mM MgSO4, 2.5 mM CaCl2, 26 mM NaHCO3, 20 mM Glucose. Adult brains slices for electrophysiological recordings were prepared as previously described (Jiang et al., 2015; Yu et al., 2012). Briefly, mice were quickly decapitated, brain were removed into ice-colded modified ASCF containing: 93 mM NMDG, 93 mM HCl, 2.5 mM KCl, 1.2 mM NaH2PO4, 30 mM NaHCO3, 20 mM HEPES, 25 mM glucose, 5 mM sodium ascorbate, 2 mM Thiourea, 3 mM sodium pyruvate, 10 mM MgSO4 and 0.5 mM CaCl2 with PH 7.35. The brain was coronally sliced to 350 μm using vibratome at a speed of 0.03 mm/s. Slices were transferred to 37°C NMDG solution to be incubated for 15 mins and then transferred to 37°C ASCF for another 1 hr. Recordings were performed in pyramidal cells within the lay II/III of mouse motor cortex under conditions of room temperature (∼24°C). Patch clamping rig is equipped with upright microscope (BX51WI, Olympus) with 60× objective lens (water-immersion, NA 1.00) and differential interference contrast (DIC). Recording pipettes (∼12 MΩ) were manufactured by a P1000 micropipette puller (Sutter Instrument, USA) and filled with the internal solution containing: 136 mM K-gluconate, 6 mM KCl, 1 mM EGTA, 2.5 Na2ATP, 10 mM HEPES (280 mOsm, pH=7.2 with KOH). Data were collected with a low-pass filter at 2 kHz using Multiclamp 700B and Digidata 1322A/D converter, and sampled at 10 kHz using pCLAMP software. Recording with series resistance > 30 mΩ were excluded for further analyses. The first action potential the elicited in response to step current injections was chosen for analyses with Clampfit (version10.7). The threshold of action potentials is documented as the membrane voltage at the first point at which the dV/dt reaches 10 V/s during the rise phase of the phase-plane plot of the membrane voltage against its change rate. Peak amplitude was calculated as the difference between threshold and peak point. MiniAnalysis (Synaptosoft, version 6.0.7) was used to document synaptic currents. The threshold was set as 8 pA for event detection.

### Confocal imaging

Immunostaining was imaged with Leica SP8 STED 3X equipped with Leica sCMOS DFC9000 at 40X (1.0 NA). Z-stack was set at 0.2-0.5 μm. The diameter of the AIS at the start, middle and end points were documented based on AnkG and βIV-spectrin immunostaining along the axon with Image J (NIH)-Fuji. The start and end of the the AIS was defined as the point of AnkG or βIV-spectrin immunoreactivity that fell below 10% of the maximum fluorescence intensity (Ko et al., 2016; Schafer et al., 2009; Zhao et al., 2020). In Emx1-Cre/COUP-TFI fl/fl mouse line and Emx1-CreER/COUP-TFI fl/fl mouse line. For conditional cKO, AISs from 5 locations containing 4 corners and 1 middle field (1/25 of the total area) in the motor cortex were chosen for analyses. Total 20 AISs from one image were averaged for each brain slice. For hippocampal AISs, total 15 AISs that were discernible from three locations (two sides and 1 middle) were analyzed for each brain slice.

For spine density, neurobiotin filling during recording was used for loss of COUP-TFI in individual neurons and Golgi staining was used for cKO mice. The dendrite length and number of spine were measured with Image J (NIH)-Fuji.

### Immunoblotting

The Immunoblotting was performed as previously described (Zhao et al., 2020). For immunoblotting, mice were anesthetized by isoflurane and quick decapitated, brain were removed and the motor cortex was dissected on the surface of an ice-colded chamber. Brain tissues were flash-frozen in liquid nitrogen and stored at -80°C until protein homogenization. Proteins were separated with the SDS-PAGE (Bis-Tris, 4-12% gradient gels) and transferred to nitrocellulose membrane at 100 mA for overnight at 4°C. Membranes were blocked with 5% BSA for 2 hrs and were incubated with a primary antibody followed by the appropriate horseradish peroxidase-conjugated secondary antibodies. The primary antibodies were used as follows: Chicken anti-Nfasc (1:500; Cat #: AF3235, R&D Systems), Mouse anti-AnkG (1:500; N106/36, NeuroMab), Rabbit anti-Nav1.6 (1:200; ASC009, Alomone). Membranes were detected by ECL detection kit (Bio-Rad). ImageJ (NIH)-Fuji software were used to quantify the relative level of proteins.

### Open filed task

Mice were placed in a square box with 50 cm wide and 45 cm high. Side walls of the open field are painted into light blue and a white board were put in the bottom of the box to increase the contrast. The box was surrounded by light tight curtain to minimize the effects of the environment on behavior tests. The box was divided into equally 4×4 square area. The central 2×2 square areas were defined as the central area and other areas were defined as the peripheral area. At the beginning of each task, mice were placed in the central area of the box. SuperMaze (Shanghai XinRuan Information Technology Co., Ltd) software was used to track the movement of mice for 5 mins video recordings.

### Elevated plus maze task

Mice were placed on an elevated plus maze consisting of two open arms and two close arms surrounded by 15 cm height opaque walls. The length of each arm is 35 cm, white boards were put in the bottom of four arms to increase contrast. Behavioral tests were conducted under an environment surrounded by a curtain. The plus maze was divided into two open arms, two close arms and a small center area. At the beginning of each task, mice were put on the center area with their heads toward an open arm. SuperMaze software were used to Track the movement of mice for 10 mins video recording.

### Virus delivery

The EGFP-Cre and EGFP pAAV2s were purchase from the company (Shanghai OBiO Technology Corp, Ltd). For virus delivery, mice were deeply anesthetized by chloral hydrate and heads were fixed in a stereotaxic apparatus, AAV2/9-CRE and EGFP viruses were injection into mouse motor cortex at the coordinates of 1.10 mm ahead of the bregma, 1.5 mm lateral to the midline and 1.0 mm from the surface. Two weeks after AAVs injection, mice were sacrificed for immunostaining and electrophysiological recordings. iCre-EGFP and COUP-TFI-IRES-EGFP were cloned into the retrovirus vector. For embryonic retrovirus delivery, pregnant mice were deeply anesthetized by isoflurane. The E14.5 mice embryo were exposed to the clean environment and retroviruses mixed with dye were injected into the lateral ventricle with glass needles. The pregnant mice were sutured and recovered.

## Statistics

Blinded data analyses to the experimental groupings were conducted by investigators. Statistical analyses and data plots were performed using the GraphPad Prism 5.0 software (Graphpad Software, Inc.). Data were tested for significance between groups using unpaired t-test or one-way / two-way ANOVA with *post hoc* Bonferroni’s multiple comparisons test, and P < 0.05 was considered significant. All figures were composed using Illustrator software (Adobe Systems, Inc.).

## Acknowledgements

We thank Dr. Song-Hai Shi (Tsinghua University, Beijing, China), Dr. Shaoyu Ge (State University of New York at Stony Brook, USA) and Dr. Guisheng Zhong (ShanghaiTech University, China) for helpful discussions. We thank Dr. Ke Tang (Nanchang University, Nanchang, China) for kindly providing the COUP-TFI fl/fl and Emx1-Cre knock-in mice and Dr. Yongchun Yu (Fudan University) for kindly providing the Emx1-CreER knock-in mice. We also thank the facilities of Imaging Core of Life School of Science and Technology at ShanghaiTech University for technical supports. This work was supported by the National Natural Science Foundation of China (31671062, 81870734) and the Shanghai Municipal Government and ShanghaiTech University.

## Author Contributions

S. H. conceived the project and designed the experiments. X. W. performed most of experiments in this study. H. L. assisted behavioral experiments. J. H. helped immunoblot experiments.

## Figures

**Supplementary Figure 1:**
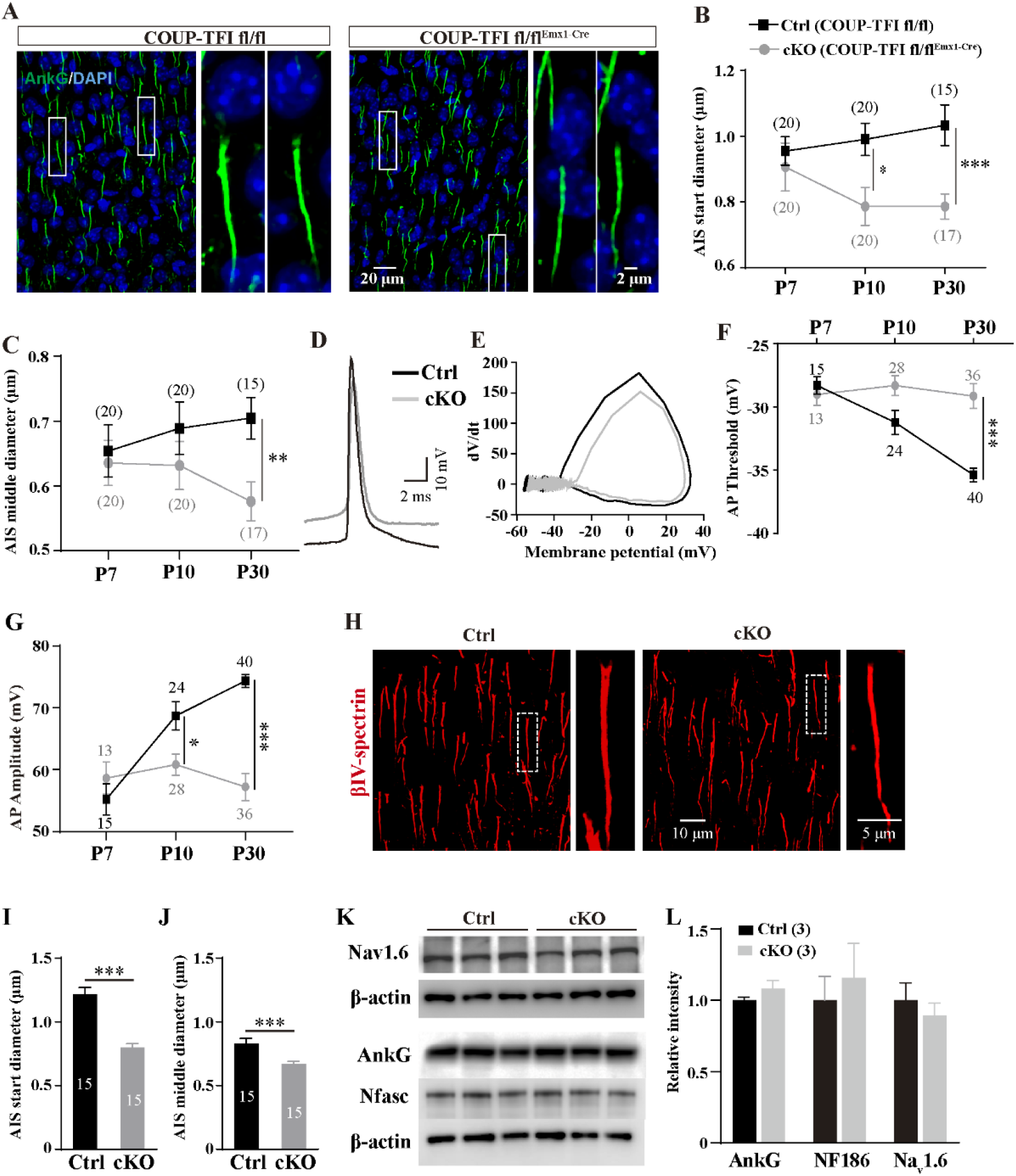
Conditional knockout of COUP-TFI reduces AIS diameter and impairs AIS functions. (A) Representative images of AnkG staining in the layer II/III of the neocortices from COUP-TFI fl/fl (control) and COUP-TFI fl/fl^Emx1-Cre^ (cKO) mice at P30. (B, C) Quantification of AIS diameter at the start point (*B*) and the middle (*C*) point for both control and cKO mice at P7, P10, and P30. (D) Overlapping of sample traces of the first APs in cortical pyramidal neurons from control and cKO mice. (E) Phase-plane plots of membrane voltage and its change rate in response to current injection in (*D*). (F, G) Quantification of AP threshold (*F*) and peak amplitude (*G*) for both control and cKO mice at P7, P10, and P30. (H) Representative images of βIV-spectrin staining in the neocortices from control and cKO mice at P30. Boxes are enlarged into the right panels. (I, J) Quantification of AIS diameter at the start point (*I*) and the middle point (*J*) bases on βIV-spectrin staining at P30 for both control and cKO mice. (K) Representative images of AnkG, Nav1.6 and neurofascin (Nfasc) immunoblots of cortical tissue homogenates from control or cKO mice. (L) Quantification showing no change in the intensity of these three protein immunoblots between controls and cKO mice. N in (B), (C), (I) and (J) represents the number of slices analyzed from 5 mice. N in (F) and (G) represents neuron number. N in (L) represents mouse number assessed. Data are presented as mean ± SEM. *, p < 0.05; **, p < 0.01; ***, p < 0.001. P value in (B), (C), (F) and (G) are determined using Two-way ANOVA with *post hoc* Bonferroni’s tests. P value in (I) and (J) are calculated using student’s t-test.

**Supplementary Figure 2:**
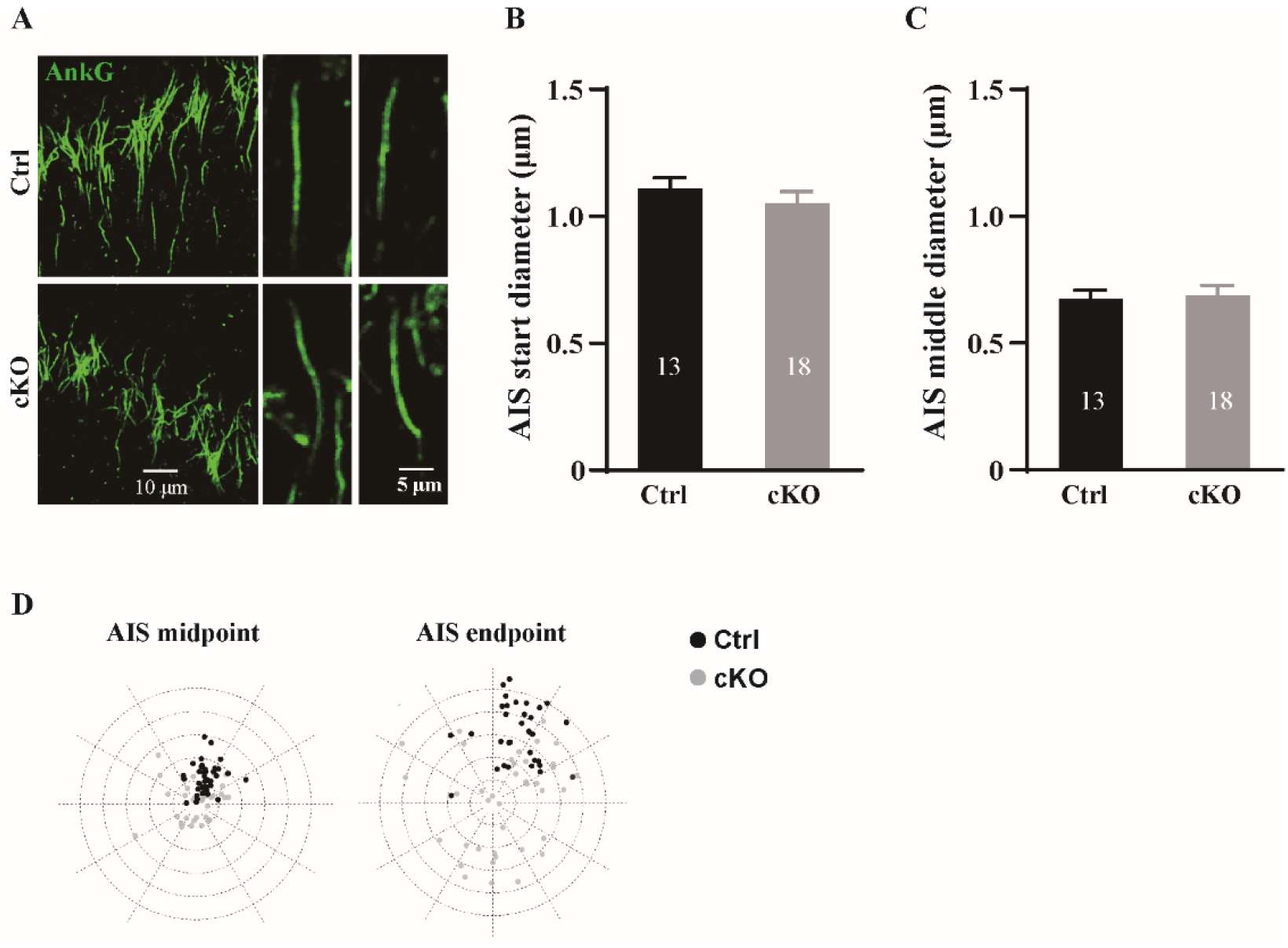
Ablation of COUP-TFI has no effect on AIS diameter in pyramidal cells of the hippocampus. (A) Representative images of AnkG staining in hippocampi from control and cKO mice at P30. (B, C) Quantification of AIS diameter at the start point (B) and the middle point (C) at P30 for both control and cKO mice. (D) Circular graph plots of the distance and direction of AIS middle point and endpoint away from the soma. The AIS tends to curve toward different directions in hippocampal pyramidal cells of cKO mice compared to controls. N represents slice number assessed from 4 Ctrl and 5 cKO mice, respectively. Data are presented as mean ± SEM.

**Supplementary Figure 3:**
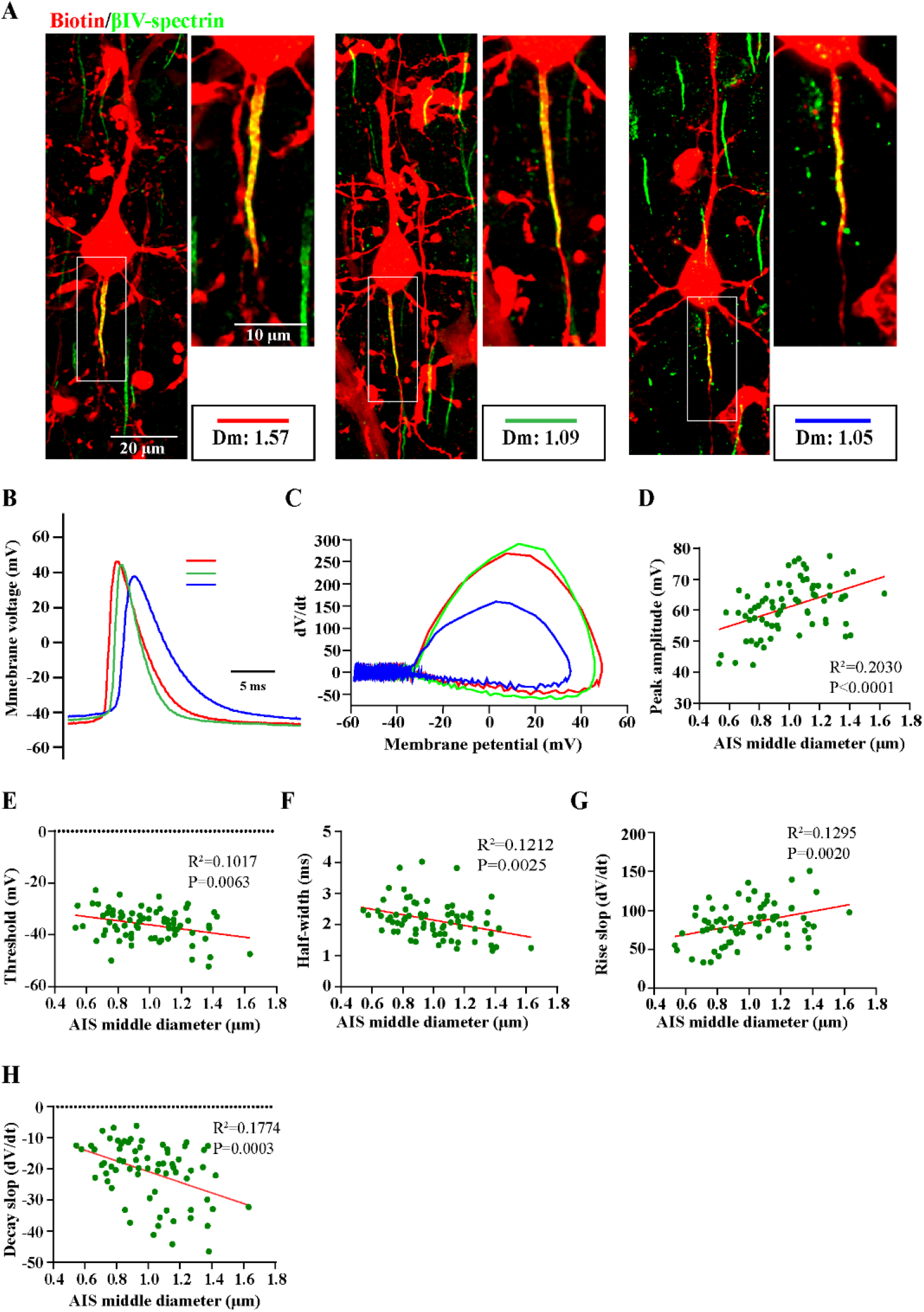
The relationship of AIS diameter and function. (A) Representative images of βIV-spectrin staining in recorded cortical pyramidal cells filled with biotin. (B) Overlapping of sample traces of the first APs from three neurons of different diameters. (C) Phase-plane plots of membrane voltage and its change rate in response to current injection from the three neurons in (*B*). (D-H) Linear plots of AIS middle diameter against AP peak amplitude (*D*), threshold (*E*), half-width (*F*), rise slop (*G*), and decay slope (*H*) in cortical pyramidal cells of the mouse neocortex. N represents neuron number assessed from 9 mice. P values are determined using t-test.

**Supplementary Figure 4:**
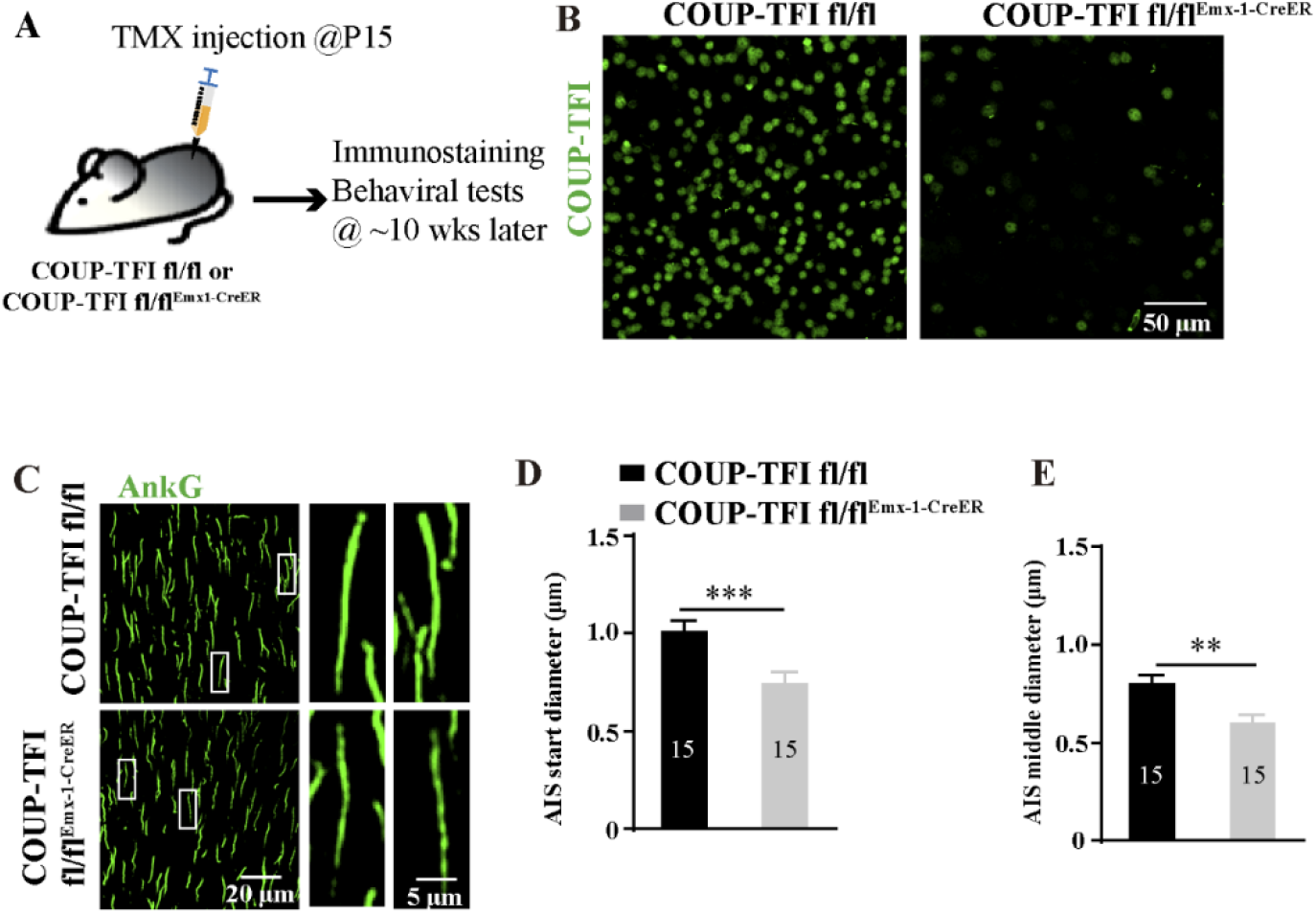
TMX-induced postnatal ablation of COUP-TFI reduced AIS diameter in the mouse cortex. (A) Schematic of the experimental paradigm for behavioral tests. Mice received intraperitoneal injection of tamoxifen at P15 and immunostaining and behavioral tests were performed ∼10 weeks later. (B, C) Representative images of COUP-TFI (*B*) and AnkG (*C*) immunostaining in the neocortices from COUP-TFI fl/fl and COUP-TFI fl/fl^Emx1-CreER^ mice. Boxes in (C) are expanded to the right panels. (D, E) Quantification of AIS diameter at the start point (*D*) and the middle (*E*) point for both control and TMX-induced cKO mice. N in (D) and (E) represents the number of slices analyzed from 5 mice. **, p < 0.01; ***, p < 0.001, student’s t-test.

